# Simple soil eDNA sampling reveals terrestrial mammal diversity in the Central Kalahari and highlights waterholes as sampling hotspots

**DOI:** 10.64898/2026.10.09.758017

**Authors:** Tamara Schenekar, Emily Bennitt, Kaisara Kelebileone, Robynne Kotze, Andrew Loveridge, Ella Mutch, Tlotlo S. Ndaba, Pooja Singh

**Affiliations:** University of Graz, Department of Biology, Graz, Austria; Okavango Research Institute, University of Botswana, Maun, Botswana; Wildlife Conservation Research Unit, Department of Biology, Oxford University, Oxford, United Kingdom. Conservation of Natural Ecosystems Trust, Maun, Botswana; Wildlife Conservation Research Unit, Department of Biology, Oxford University, Oxford, United Kingdom, Panthera, New York, USA; Wildlife Conservation Research Unit, Department of Biology, Oxford University, Oxford, United Kingdom; Division of Aquatic Ecology and Evolution, Institute of Ecology and Evolution, University of Bern, Bern 3012, Switzerland; Department of Fish Ecology and Evolution, Swiss Federal Institute of Aquatic Science and Technology (EAWAG), Kastani

**Keywords:** DNA metabarcoding, soil, terrestrial mammals, Central Kalahari Game Reserve, camera trapping, biodiversity monitoring, protected areas

## Abstract

Remote and vast ecosystems present a challenge for biodiversity monitoring. The Central Kalahari Game Reserve in Botswana is one such ecosystem spanning 52,800 km^2^, a mosaic of fossilized rivers, clay pans, semi-arid sand dunes, scrub, and woodland. It is one of the last remaining untouched wildernesses in the world and hosts unique biodiversity, including populations of threatened mammals. Due to its extreme arid environment, size, and inaccessibility, surveying biodiversity across the reserve is both logistically and technically difficult. Here we demonstrate that a low-effort approach of sampling soil across the reserve is an efficient way to rapidly assess the terrestrial mammal biodiversity using eDNA metabarcoding. We found that eDNA from soil near waterholes performed better at capturing local biodiversity than soil from roads. eDNA metabarcoding detected 38 mammal species, 15 bird species and two amphibians and a simultaneously conducted camera trapping study detected 29 mammal species. eDNA was particularly powerful for detecting small-bodied taxa, including murids, mongooses and bats, which were largely missed by camera traps, while camera trapping was aimed at and performed well for detecting felids. Mammal species that had a higher activity nearby eDNA sampling points (i.e., a higher number of camera trapping recordings nearby) had a higher probability of being detected via eDNA. We propose that simple soil collection for eDNA metabarcoding, particularly close to animal congregation points such as savanna waterholes, offers a practical, scalable and cost-effective strategy for DNA-based biomonitoring in large, remote areas such those in Southern Africa.

## Introduction

Biodiversity monitoring is a key tool for quantifying and ultimately mitigating ongoing global biodiversity loss (Pereira and David Cooper 2006). Yet, current monitoring efforts are markedly uneven across regions, with a disproportionate focus on North America, Europe and Australia, while much of Africa, South America and South-East Asia remains sparsely monitored (Di Marco et al. 2017; Schmeller et al. 2017). This geographic bias is coupled with a disconnect between the monitoring focus and conservation needs, as the data-poor regions are often rich in biodiversity and subject to the most severe threats to biodiversity (Collen et al. 2008; Schmeller et al. 2017). For example, the largest areas experiencing the greatest numbers of mammal species extinctions are located in Africa and Southeast Asia (Pereira and David Cooper 2006). The reasons for this imbalance are associated with socio-economic legacies rooted in colonial history that persist in present day GDP (Waldock et al. 2026) and affect governmental monitoring. Lower-middle and low-income countries, which include many of the world’s megadiverse regions especially along the tropics, typically have the least governmental support for biomonitoring programs (Moussy et al. 2022). In addition, logistical challenges arising from limited infrastructure, such as sparse road networks, inadequate storage facilities for field equipment, restricted access to laboratory space, and the need to transport bulky survey gear over long distances to remote areas, further complicate biodiversity monitoring in any wild and remote area (Morse et al. 2007; Mezgebu 2020; Penhacek et al. 2025). Hence, ironically, the very inaccessibility that often preserved biodiversity in these areas initially, now frequently hampers the assessment of biodiversity threats in these regions, complicating conservation efforts.

Savannas, semi-arid landscapes in Africa that are characterized by a mix between grass- and woodlands, are one of the most biodiverse ecosystems supporting large mammals, with ungulate biodiversity and herbivore biomasses being among the highest ever quantified (Bourliere and Hadley 1970; McNaughton and Georgiadis 1986). These vast landscapes cover 50% of the African continent and suffer from several anthropogenic pressures such as land cover change, landscape fragmentation and climate change (Osborne et al. 2018). Habitat degradation presents a severe threat to resident wildlife, such as the iconic large herds of dry-adapted migratory herbivores (Bolger et al. 2007) and, in turn, the charismatic African carnivore populations that depend on them, for which protected areas often constitute the last remaining strongholds (Bodasing 2022).

The Central Kalahari Game Reserve (CKGR) lies in the heart of the Kalahari Desert, a semi-arid savanna system spanning parts of Namibia, Botswana and South Africa. The CKGR is located in central Botswana and covers over 52,800 km², making it the largest terrestrial protected area in southern Africa and the second largest game reserve worldwide, exceeding the land area of several sovereign states, including Slovakia and Costa Rica.

The establishment of the park by the Botswana government and the low impact of indigenous peoples and their traditional ways of life has helped to maintain a rich mammal diversity in the CKGR (Perkins 2019). Large herds of migratory and dry-adapted antelopes like hartebeest (*Alcelaphus buselaphus*), blue wildebeest (*Connochaetes taurinus*), springbok (*Antidorcas marsupialis*), eland (*Tragelaphus oryx)*, and gemsbok (*Oryx gazella*), are found in the reserve, as well as carnivores including lion (*Panthera leo*), leopard (*Panthera pardus*), cheetah (*Acinonyx jubatus*), wild dog (*Lycaon pictus*), spotted hyaena (*Crocuta crocuta*), brown hyaena (*Parahyaena brunnea*) and black-backed jackal (*Lupulella mesomelas*) (Dawson and Butynski 1975; Selebatso et al. 2018a, b). Overall, the GBIF (Global Biodiversity Information Facility) database lists 52 mammal species that have been directly observed within the CKGR area (GBIF.org 2026). Small mammal diversity is presumably underestimated by this dataset. The reserve supports a diverse avifauna of almost 350 recorded species (GBIF.org 2026). Characteristic species include large terrestrial birds such as ostrich (*Struthio camelus)* and the national bird of Botswana – kori bustard (*Ardeotis kori*), alongside characteristic Kalahari specialists like Burchell’s sandgrouse *(Pterocles burchelli*) and birds of prey including pale chanting goshawk (*Melierax canorus*), tawny eagle (*Aquila rapax*), bateleur (*Terathopius ecaudatus*), secretary bird (*Sagittarius serpentarius)* and several vulture species. Furthermore, the reserve hosts a diverse assemblage of granivorous passerines. Information on amphibians and reptiles in the CKGR is comparatively limited, and the herpetofauna remains less well documented than the region’s birds and mammals. While amphibian occurrence is strongly constrained by the semi-arid climate and largely restricted to seasonal, rain-filled depressions and artificial waterholes, the reserve is expected to support a suite of arid-adapted reptiles typical of the wider Kalahari basin, including geckos (Gekkonidae), skinks (Scincidae), agamas (Agamidae), and a variety of snakes typical of Kalahari dune systems (Haake 1984).

Wildlife monitoring in the CKGR has to date relied on a limited set of approaches. For ungulate populations aerial surveys are periodically conducted (e.g., DWNP 2012). On-the-ground surveys in the semi-arid Kalahari Desert include track surveys combined with GPS collaring and indigenous knowledge (Gielen et al. 2024; Finerty et al. 2024). Furthermore, camera trapping surveys have been employed in the area, mostly by non-governmental organisations targeting large carnivores. However, in general, biodiversity assessments have been comparatively sparse. Key reasons for this are the extreme logistical challenges associated with the CKGR’s size and remoteness. Due to the vastness and relatively low vegetation productivity of the site, wildlife populations exhibit relatively low population densities and extensive home ranges, complicating monitoring even further (Finerty et al. 2024).

Environmental DNA (eDNA)-based approaches have been developed as powerful tools for biodiversity monitoring (Bohmann et al. 2014; Cristescu and Hebert 2018; Beng and Corlett 2020). Environmental DNA is DNA captured from an environmental sample without pre-isolating any target organisms (Taberlet et al. 2012). Such environmental samples can come from any source such as water, soil, sediments, air, gut contents, pollen, and more. Environmental DNA metabarcoding uses High-Throughput-Sequencing (HTS) to simultaneously sequence previously amplified metabarcoding markers in an eDNA sample in order to identify many taxa in parallel. With metabarcoding data, inferences on species composition can be made rapidly and cost-efficiently (Deiner et al. 2017). Due to the abovementioned socioeconomic constraints, there is also a heavy geographic bias in eDNA research, with most studies being conducted in temperate systems in the Global North (Belle et al. 2019; Schenekar 2023). Furthermore, freshwater and marine systems far outnumber terrestrial ecosystems in eDNA-based studies (Sahu et al. 2025), resulting in most methodological advances and interpretative frameworks being developed in aquatic contexts. Although eDNA sampling is often portrayed as an easy-to-use approach, it requires strict adherence to best practices (Goldberg et al. 2016; Wilcox et al. 2018) and is highly susceptible to contamination because it targets very low quantities of DNA (Sepulveda et al. 2020; Hutchins et al. 2022). This necessitates clean laboratory space for sample processing. However, such facilities are rarely available in remote regions. This makes it imperative to develop and test streamlined sampling protocols that minimize sample handling and reliance on bulky equipment when implementing eDNA-based monitoring remote and difficult to access parts of the planet (Schenekar et al. 2024).

In semi-arid savanna ecosystems, there are two main landscape features that present themselves as particularly suitable for eDNA-based monitoring: 1) waterholes, which are key aggregation points for wildlife in otherwise dry habitats and thus represent high-value focal sites for biodiversity monitoring (Redfern et al. 2005; Sutherland et al. 2018; Schenekar et al. 2026) and 2) road tracks, that often function as primary travel corridors for a wide range of mammal species (Abrahms et al. 2016; Fitzpatrick et al. 2024). In contrast to waterholes, where equipment is more prone to damage through wildlife due to prolonged animal presence and increased interactions with equipment, camera traps are typically deployed along wildlife trails or car tracks, as these locations offer both high animal encounter rates and a lower risk of equipment loss.

This study aimed to test the efficacy of an eDNA-based approach to monitoring biodiversity in a highly remote wilderness area. We apply eDNA metabarcoding for the first time to assess vertebrate biodiversity in the Central Kalahari Game Reserve (CKGR) in Botswana. Our main objectives were as follows: 1) To test the performance of a simple, field-adapted sampling protocol for soil eDNA, 2) to compare the efficiency of a road transect-based approach to a waterhole-focused sampling, 3) to investigate potential spatial patterns in the detected biodiversity, 4) to benchmark the sensitivity of soil-sourced eDNA metabarcoding against the known vertebrate biodiversity in the reserve with a particular focus on terrestrial mammals, and 5) evaluate eDNA-based mammal biomonitoring against camera trapping data.

## Methods

### Study site and eDNA sampling

The study site is situated in the northern part of the CKGR. The roads within the CKGR are generally limited to a very sparse network of unpaved and often narrow sand tracks that provide access to a small number of campsites and borehole-fed waterholes. These narrow tracks are often deeply sandy, and seasonally impacted by rain, making large parts of the reserve effectively inaccessible, especially during the wet season. Navigating the area requires a high-clearance 4×4 vehicle and conditions only allow short distances to be travelled per day. No access to potable freshwater, electricity, fuel or food is available within the reserve. The climate of the CKGR area is subtropical and semi-arid, with most of the rainfall occurring in the wet season from October to April. Extreme temperatures ranging from −6°C in winter to 43°C in summer have been reported (DHV 1980; Williamson and Williamson 1985). Ecologically, the area is classified as Kalahari semi-desert, Kalahari savanna or Kalahari sandveld, characterized by deep aeolian sands largely covered by extensive grasslands and interspersed with shrubs and thorn trees, including species of *Vachellia* and *Senegalia* (Sayre et al. 2013). The predominantly flat to gently rolling landscape is intersected by ancient, fossilized river valleys and associated clay-rich depressions that accumulate water after rainfall and form ephemeral pans and salt pans. Within the reserve, there is no permanent natural surface water. Freshwater is provided at ten solar panel powered artificial waterholes, serving as crucial water sources for the resident wildlife in the reserve (Selebatso et al. 2018a).

Environmental DNA sampling took place between Matswere and Tsau Gate in September 17^th^-19^th^, 2024, along the sand tracks that provide access to the selected waterholes for sampling. (Fig. 1).

**Fig. 1.**
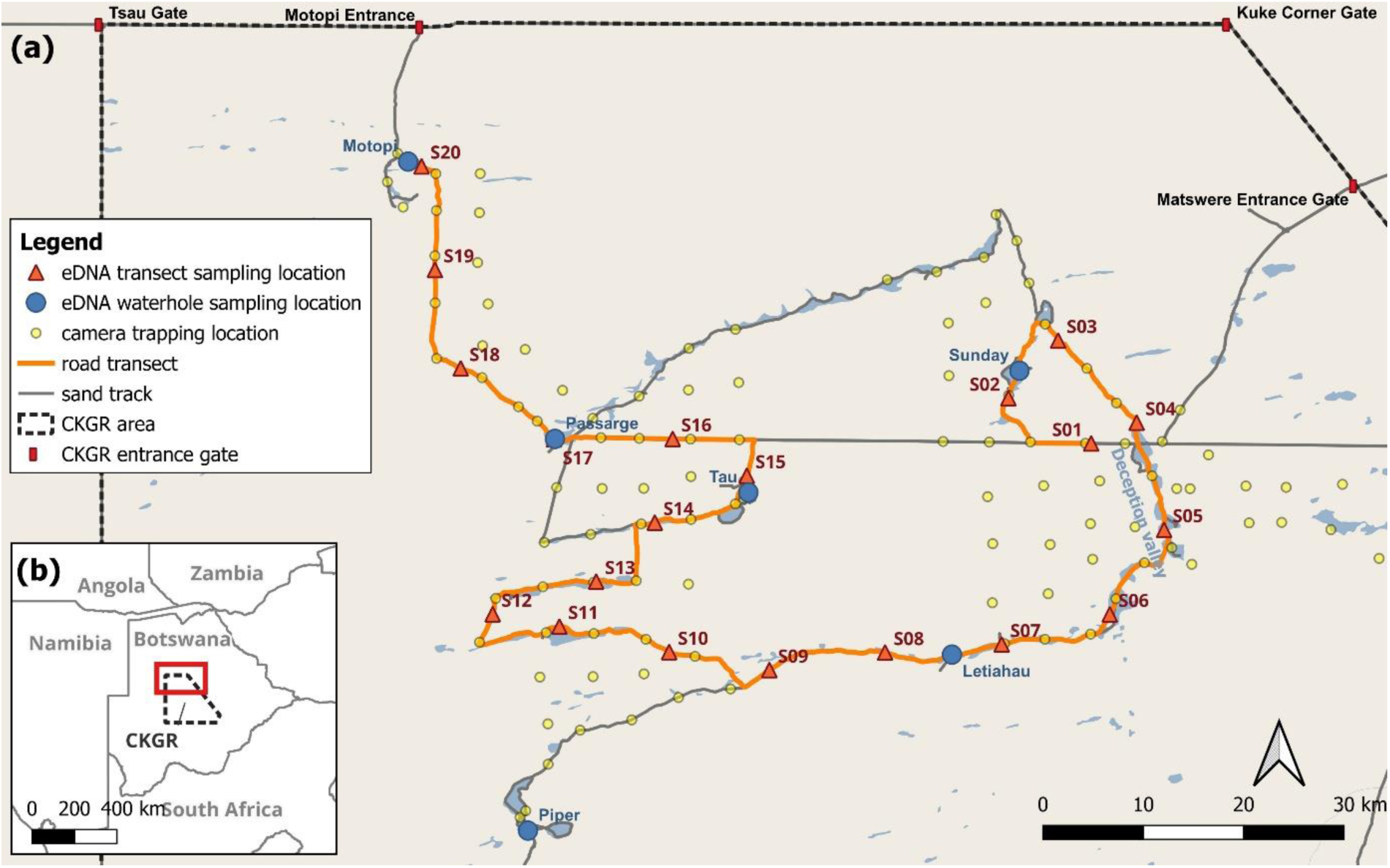
Environmental DNA sampling locations in the Central Kalahari Game Reserve. Labelled blue circles represent waterhole sampling locations, while orange triangles represent sampling locations along the road transect (orange line). Semitransparent yellow circles indicate positions of camera traps. Blue shadings in the basemap represent salt pans. Basemap data © OpenStreetMap contributors, available under the Open Database EnvLicense (ODbL).

The road transects consisted of 20 evenly spaced sampling sites along the driven sand roads used to access the selected waterholes for sampling. The distance between the transect sampling locations measured 12.235 km each and the total transect length was 234.616 km. In addition, we sampled five waterholes (Sunday, Letiahau, Tau, Passarge and Motopi) by collecting eDNA at four sites along the surrounding sand tracks, evenly spaced around each waterhole and located within 100 m of the water body, resulting in 20 waterhole-focused sampling sites. At each site (road transect and waterhole), two soil samples were collected by scooping approximately 3 ml of sand (one from the left and one from the right side of the road) into sterile 15 ml centrifuge tubes. At the end of each sampling day (within up to 12 hours of collection), 6 ml of Longmire lysis buffer (Longmire et al. 1997) was added to each sample using a micropipette. Within four days, the tubes were transported at ambient temperature and subsequently stored at −20°C at the Okavango Research Institute of the University of Botswana in Maun. On each sampling day, one field negative control was processed by filling an empty 15 ml centrifuge tube with 6 ml of Longmire lysis buffer. Gloves were worn during sample collection and the addition of Longmire buffer to minimize contamination. One waterhole (Motopi) was dry at the time of sampling. However, soil samples were collected at this location in the same manner as for the other waterholes.

### DNA extraction, metabarcoding library preparation and sequencing

All laboratory sampling processing was conducted at the Institute of Biology, University of Graz, Austria. All DNA extractions and PCR setups were performed in a dedicated low-template DNA facility, using separate laminar-flow hoods for extraction and PCR preparation to reduce contamination risk. Each extraction batch (up to 11 samples per batch) included one extraction blank, and the laminar-flow work benches were decontaminated between sessions by 30 min UV irradiation. DNA was extracted with the DNeasy PowerSoil Pro Kit (Qiagen), following the protocol of Schenekar et al. (2024) and eluted in a final volume of 100 µl. All DNA extracts were treated with the OneStep Inhibitor Removal Kit (Zymo Research) according to the manufacturer’s instructions. For metabarcoding, we used two primer sets targeting distinct fragments of the mitochondrial 12S rRNA gene: (1) the MiMammal which predominantly target mammals, and (2) the 12SV5 primer set designed for vertebrate detection (Table 1). Both primer sets carried 8-bp indices at the 5′ ends of forward and reverse primers to allow sample demultiplexing via unique dual indices.

**Table 1.**
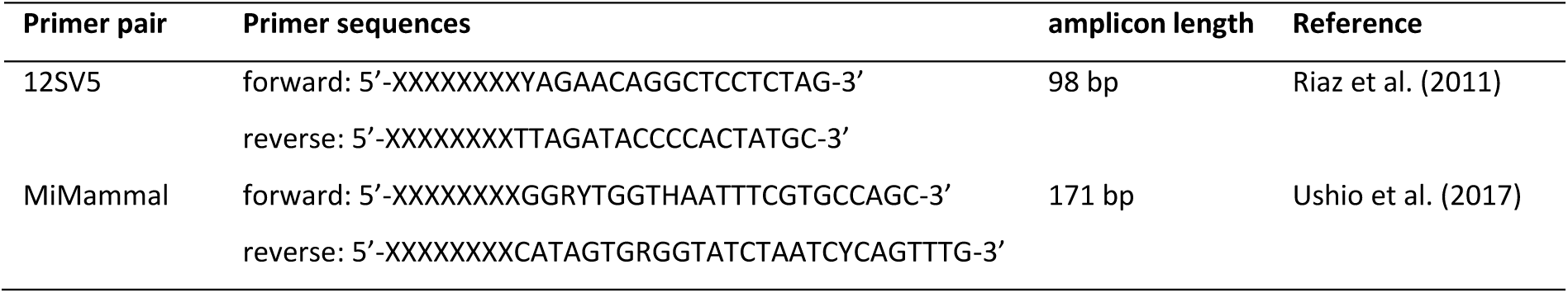
Details on the utilized primer pairs for metabarcoding in this study. XXXXXXXX indicate the 8-bp indices.

For PCR amplification four PCR replicates were conducted per sample and primer pair. Each PCR reaction consisted of 2.5 µl template DNA, 6.25 µl TaqMan Environmental Master Mix 2.0 (Thermo Fisher Scientific) 0.6 µM of forward and reverse primer, respectively and nuclease-free water up to final reaction volume of 12.5 µl. Cycling conditions were as follows: Initial denaturation at 95°C for 30 sec, followed by 40 cycles of denaturation at 95°C for 30 sec, annealing at 60°C (MiMammal) or 57°C (12SV5) for 30 sec, extension at 60°C for 30 sec and a final extension at 72°C for 7 min. All PCR products were excised from a 2% agarose gel whereby the four PCR replicates of each sample and primer pair were pooled, and subsequently cleaned using the Monarch Spin DNA Gel Extraction Kit (New England Biolabs). Four PCR positive controls, consisting of the DNA extracts of six Eurasian mammals (Eurasian beaver -*Castor fiber*, European hedgehog – *Erinaceus europaeus,* raccoon -*Procyon lotor*, brown bear -*Ursus arctos*, Eurasian fish otter – *Lutra lutra* and roe deer -*Capreolus capreolus* at a final concentration of 7 ng/µl) and two PCR negative controls (HPLC-purified water) were processed along with the samples. All controls (field-, extraction- and PCR-negative controls as well as PCR-positive control) were processed in the same manner as field samples. All 192 amplicons (two markers each from 80 field samples, three field blanks, seven extraction blanks, four PCR positive controls and two PCR negative controls) were pooled, concentrated using the DNA Clean and Concentrator-5 Kit (Zymo Research) and sent to a commercial Illumina sequencing provider for final library preparation (using the NEBNext® Ultra™ II DNA PCR-free Library Prep Kit for Illumina, New England Biolabs) and sequencing. Libraries were sequenced on an Illumina NovaSeq X in 150 bp paired-end read mode with a targeted output of 30 Gigabases (i.e., approx. 521,000 reads per sample and marker).

### Bioinformatic analysis

After checking quality of the raw reads via FastQC (Andrews 2010), quality filtering was conducted via *vsearch* v2.15.1 (Rognes et al. 2016) using the *fastx_filter* function (error rate: fastq_maxee = 1). Read pairs were merged using the *fastq_mergepairs* function, allowing for merging of staggered reads and using default settings otherwise. Samples were demultiplexed using *cutadapt* 4.1 (Martin 2011), by indices (allowing for one error) and the by primer (allowing for two errors), followed by length filtering (filtered length for 12SV5: 85 – 115 bp; MiMammal: 155-190 bp). Dereplication and creation of amplicon sequence variants (ASVs) were conducted with the *derep_fulllength* function and only ASVs that had a minimum read count of five reads and a minimum alpha of 2 were retained using the *unoise_alpha* algorithm. Chimeric sequences were removed using the *uchime3_denovo* function. Reads were mapped to ASVs with the *usearch_global* function using a threshold of 99% similarity. Taxonomic assignment was done against a custom database created from the srRNA_SINTAX database of MIDORI2 (v. GB268; Leray et al., 2022). For database curation, a species list of all previously documented vertebrates in Botswana was retrieved querying the GBIF database for with the following filters: TaxonKey = Chordata, Country = Botswana, OccurrenceStatus = Present, BasisOfRecord = Machine Observation or Human Observation or Occurrence evidence (GBIF.org 2026). From these species names, the corresponding taxonomic identifiers (taxIDs) were retrieved using NCBI’s TaxIdentifier tool (NCBI 2020) while carefully checking for potential mismatches in nomenclature between databases due to synonyms. The MIDORI2 database was then filtered for these taxIDs, also excluding sequences with ambiguous bases using a custom bash script. This database was then queried for the ASV sequences with the *sintax* algorithm of *vsearch* and a sintax_cutoff of 0.9. Only ASVs that could be assigned to the species level were retained for analysis and ASV assigned to the same species were amalgamated. Read counts were transformed to binary detection/non-detection data only. For this, we applied a dynamic detection threshold based on read counts in the control samples (field-extraction- and PCR negative controls and PCR positive controls): For a given sample and species, a detection was only counted as a true detection if the read counts in the given sample exceeded the read counts in any of the control samples.

The resulting list of detected species was then carefully screened for ecologically implausible occurrences. Based on this, the following species were excluded from further analysis: (1) All domestic or human-associated animals: cattle (*Bos taurus*), domestic dog (*Canis lupus*), domestic goat (*Capra hircus*), rat (*Rattus norvegicus*) pig (*Sus scrofa*) (2) Black rhinoceros (*Diceros bicornis*): No black rhinoceros have been reported from CKGR. Furthermore, the SINTAX score of (0.90) of the ASV assigned to this species falls just at the confidence threshold applied. Therefore a taxonomic misclassification of a resident odd-toed ungulate or secondary DNA input into the reserve through vehicle tires or vultures, as described in Schenekar et al. (2026) are likely sources of this detection. (3) Waterbuck (*Kobus ellipsiprymnus*): This species is highly water dependent and has not been reported within the reserve before, this detection is also assumed to stem from secondary DNA input. (4) Sable antelope (*Hippotragus niger*): This species is relatively water dependent as well, occurs in more mesic savanna systems and has not been documented before. In contrast, the congeneric roan antelope (*Hippotragus equinus*) can tolerate drier conditions than sable and has been documented in the CKGR before. Only detected by the shorter 12SV5 fragment, two of the three ASVs assigned to sable have an almost equal mean genetic distance to sable and roan (6.8 bp versus 7.0 bp and 7.8 bp versus 8.0 bp) with the third ASV being closer associated with sable (0.8 bp versus 4.0 bp) but having only very few informative sites. Due to the ecological plausibility, we therefore conclude this to be a taxonomic misassignment and re-assigned sable detections to roan detections.

### Camera trapping

Between September 4^th^ and November 9^th^, 2024, two camera trap surveys were set up across the top third of the CKGR. The eastern survey and western survey consisted of 61 and 62 camera trap stations respectively (Fig. 1). Surveys were designed in such a way as to maximize recaptures of large carnivores such as lion and leopard and consisted of a grid layout with stations between 4 and 5 km apart (du Preez et al. 2014). The grid was designed prior to survey deployment using Google Earth, to ensure optimal trap spacing. Once in the field, camera stations were placed within 500 m of the original point. As large carnivores commonly use roads and game trails when traversing protected areas (Cusack et al. 2015), camera placement was prioritized on roads as far as possible, with some stations placed offroad on obvious game trails to maintain intactness of the gird, and at waterholes to further maximize large carnivore captures.

Each camera station consisted of two xenon-flash V7 PantheraCams (Panthera, New York, USA), approximately 8 meters apart, and slightly offset to avoid flash interference. Cameras were secured to 1.2 m high metal stakes hammered into the ground, with the cameras at a height of approximately 60 cm above the ground. Vegetation was cleared for approximately 2 m radius around each stake and in front of the cameras to reduce triggers caused by vegetation. The flash was set to 3 m, with cameras set to take a two-image burst approximately 15 seconds apart when triggered by movement. Once the cameras were set up, details pertaining to the station including GPS coordinates, camera models and numbers, and vegetation type were recorded, along with time and date of setup. Camera stations were checked at least twice during the survey to ensure cameras did not have low batteries and did not suffer interference from wildlife. Once the survey was completed, data were downloaded onto hard drives, and uploaded onto Traptagger (traptagger.co.uk), an AI-based species identification platform. Data upload included an upload of trap locations, to ensure that each image was associated with the correct geographic location. Once data was sorted to species level through a combination of AI and manual sorting, species data was quality checked to reduce chances of misidentification.

### Statistical analysis

Statistical analysis was conducted in R 4.4.1 (R Core Team 2023) via RStudio (Posit team 2024). Alpha diversity derived from eDNA data was calculated as species richness and summarized by location type as well as overall and per-site. Overlap and uniqueness in the detected species between location type was visualized in an area-proportional Euler diagram using the eulerr package (Larsson 2024). We compared species richness between waterhole and transect sites using a generalized linear modelling framework. The number of species detected per site was modelled as a count response with location type (waterhole vs. road transect) as the explanatory factor. We initially fitted a Poisson generalized linear model and evaluated model fit by inspecting the Pearson-based dispersion statistic, which indicated substantial overdispersion. We then refitted the model using a negative binomial error distribution with a log link using the function glm.nb in the MASS package (Venables and Ripley 2002). Model assumptions and residual structure were assessed using simulation-based diagnostic procedures implemented in DHARMa (Hartig 2024), including tests for overdispersion and zero inflation. The effect of location type on species richness was then evaluated by likelihood ratio tests (Chi-squared) comparing models with and without the location type term. To examine differences in beta diversity between waterhole and transect sites, we computed a Jaccard dissimilarity matrix using the vegdist() function in vegan (Oksanen et al. 2025) and differences in community composition between site types were tested using a permutational multivariate analysis of variance (PERMANOVA) implemented in vegan with location type as the grouping factor. We also visualized patterns in community composition among sites using non-metric multidimensional scaling (NMDS) based on Jaccard dissimilarity, fitted with metaMDS () in vegan, plotted with ggplot2.

We assessed whether community similarity among road transect eDNA sampling sites declined with increasing geographic distance (isolation by distance) using a Mantel test. Pairwise community dissimilarities among the road transect sites were calculated using Jaccard distance with the function vegdist() in vegan. In parallel, geographic distances (distance travelled along the transect) among road transect eDNA sampling sites were converted to a distance matrix and Mantel test was conducted via mantel() in vegan, using Pearson correlation between the community and geographic distance matrices and 9 999 permutations to assess significance.

In order to benchmark the sensitivity of the eDNA survey to capture ambient vertebrate biodiversity against to documented (and publicly available) biodiversity information, we compared the detected species list to the species occurrence data in GBIF database. For this we queried GBIF with the following filters: Scientific name = Chordata, Occurrence status = present, Basis of record = Human observation or Machine observation, Has coordinates = True, Has geospatial issue = false. Geometry: multipolygon of the CKGR area. Dataset of the query: GBIF.org (2026). The resulting species list was then compared to the detected species of the eDNA survey.

To compare the detected mammalian biodiversity with the simultaneously conducted camera-trapping survey, a comprehensive species list of all taxa recorded by all camera traps across both surveys (east and west, later on referred to as one camera trap survey) was compiled. Only mammal taxa that could be reliably identified to species level were retained for subsequent analyses. Based on these data, we generated lists of species detected exclusively by eDNA, exclusively by camera trapping, and those detected by both methods.

To assess whether mammals that were more frequently detected by camera-trapping had a higher chance of eDNA detection, we applied a binary logistic regression. Specifically, we modelled eDNA detection (presence/absence per species) as a function of the total number of camera-trap image detections per species, using a binomial error distribution with a logit link. Additionally, to assess whether spatial information on the camera-trap detection had an effect on a species’ detection at a given eDNA sampling site, we computed a distance-weighted camera activity index for each eDNA sampling site and species. For this, we computed a distance weight for each camera trap site and eDNA site combination by applying an exponential decay function: w(d) = e^(-d/λ)^ with d being the air distance between eDNA sampling site and camera trap site and λ = 5000 m. The chosen λ is expected to reflect an ecologically plausible scale at which most relevant camera trap activity occurs. The number of camera trap images for each species at a given camera trapping site was multiplied by w and for each eDNA site and species, this weighted image count was summed over all camera trapping sites to receive respective camera activity index. This index was then used as a predictor in a binomial GLMM of eDNA detection (presence/absence) with random intercepts for site and species using the lme4 (Bates et al. 2015) R package. All R codes and input files for statistical analysis can be found in Supporting Information 1.

## Results

### Metabarcoding sequencing data

A total of 68.5 Gbases (228,388,192 sequence read pairs) of raw data were generated. Of these, 174,026,559 (76.2%) reads remained after quality filtering, pair merging and demultiplexing – 102,852,361 for 12SV5 (approx. 1,189,500 reads per sample) and 71,174,198 for MiMammal (approx. 741,400 reads per sample). A total of 730 and 1220 ASVs were identified for 12SV5 and MiMammal, respectively. To these, 92.25% (12SV5) and 91.44% (MiMammal) of the reads could be mapped. Sequencing data can be accessed through NCBI’s GenBank (BioProject ID PRJNA1524099).

### Alpha and beta diversity

A total of 55 vertebrate species were detected by the eDNA survey, of which 38 were mammals, 15 were birds and two were amphibians (Table 2, Supplementary Table S1). No reptiles were detected with this survey. Of the 55 species detected, 27 were detected by both waterhole and road transect sampling approaches. Twenty-six species were detected only at waterhole sites and two species detected only at road transect sites (Fig. 2).

**Fig. 2.**
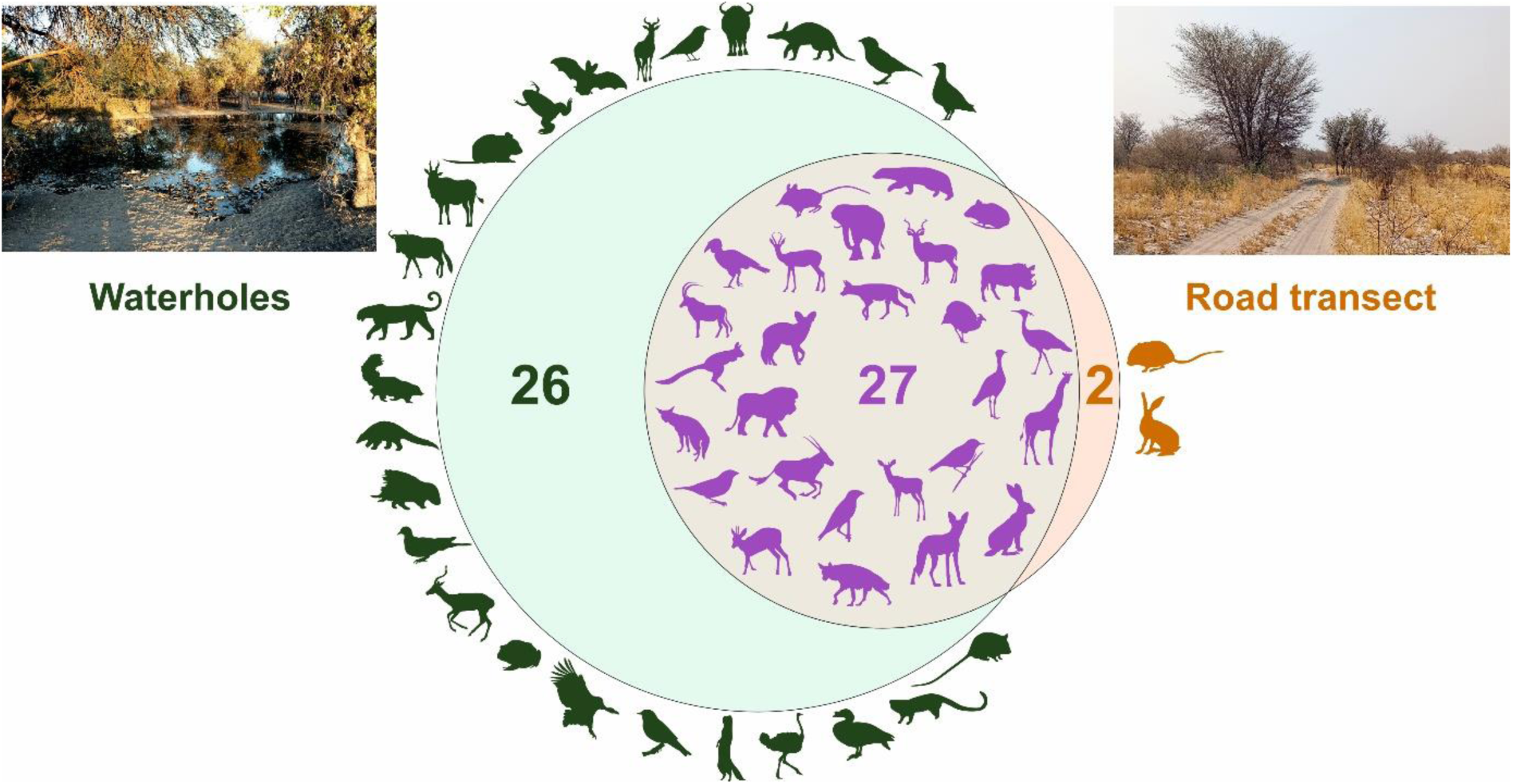
Overlap of detected species between the two sampling approaches of soil-derived eDNA metabarcoding survey in Central Kalahari Game Reserve. Species detected exclusively from the waterhole-focused sampling approach are depicted on the left in green, those from the road-transect sampling approach on the right in orange. Species detected by both sampling approaches are shown in the center in purple.

**Table 2.**
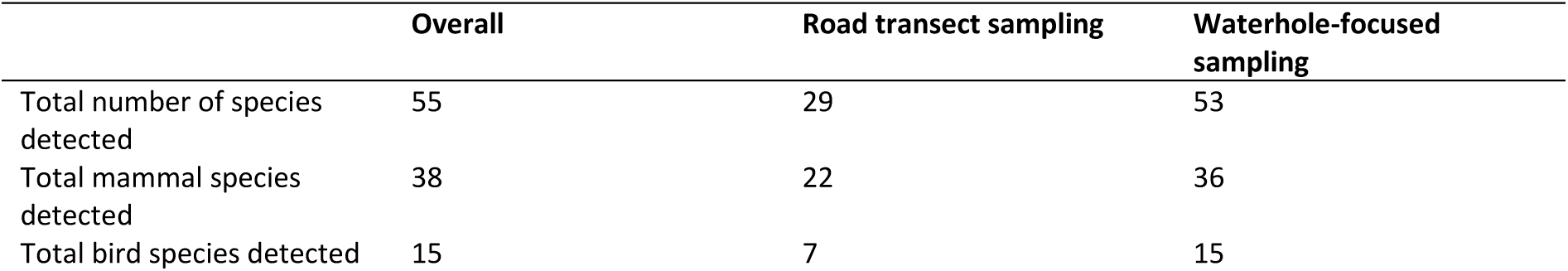
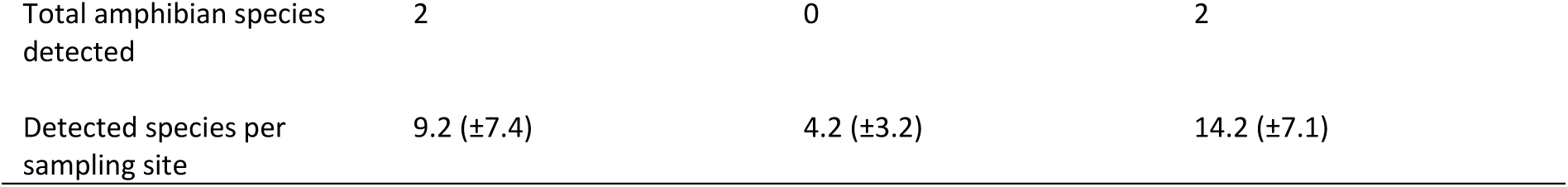
Detected alpha diversity (species richness) in the eDNA survey, shown overall and separately for the two sampling approaches (waterhole and road transect). For per-site values, mean values with standard deviations in brackets are given.

|  | Overall | Road transect sampling | Waterhole-focused sampling |
| --- | --- | --- | --- |
| Total number of species detected | 55 | 29 | 53 |
| Total mammal species detected | 38 | 22 | 36 |
| Total bird species detected | 15 | 7 | 15 |
| Total amphibian species detected | 2 | 0 | 2 |
| Detected species per sampling site | 9.2 ($\pm 7.4$) | 4.2 ($\pm 3.2$) | 14.2 ($\pm 7.1$) |

We found that the number of species detected per site was strongly influenced by location type (road transect versus waterhole). In the negative binomial GLM, species richness was significantly higher at waterholes than along road transects (Table 3, Fig. 3a, Fig.4). The model estimated that waterhole sites had, on average, 3.4 times more detected species than transect sites. Model diagnostics based on DHARMa residual simulations indicated no remaining overdispersion (p = 0.80) and no evidence for zero inflation (p = 0.72). The likelihood-ratio test indicated a better model fit when including the location type parameter (χ^2^ = 36.63, df = 38, p < 0.001). In addition to a lower per-sample species diversity, road samples were more heterogeneous in species composition across samples (Fig. 3b).

**Figure 3.**
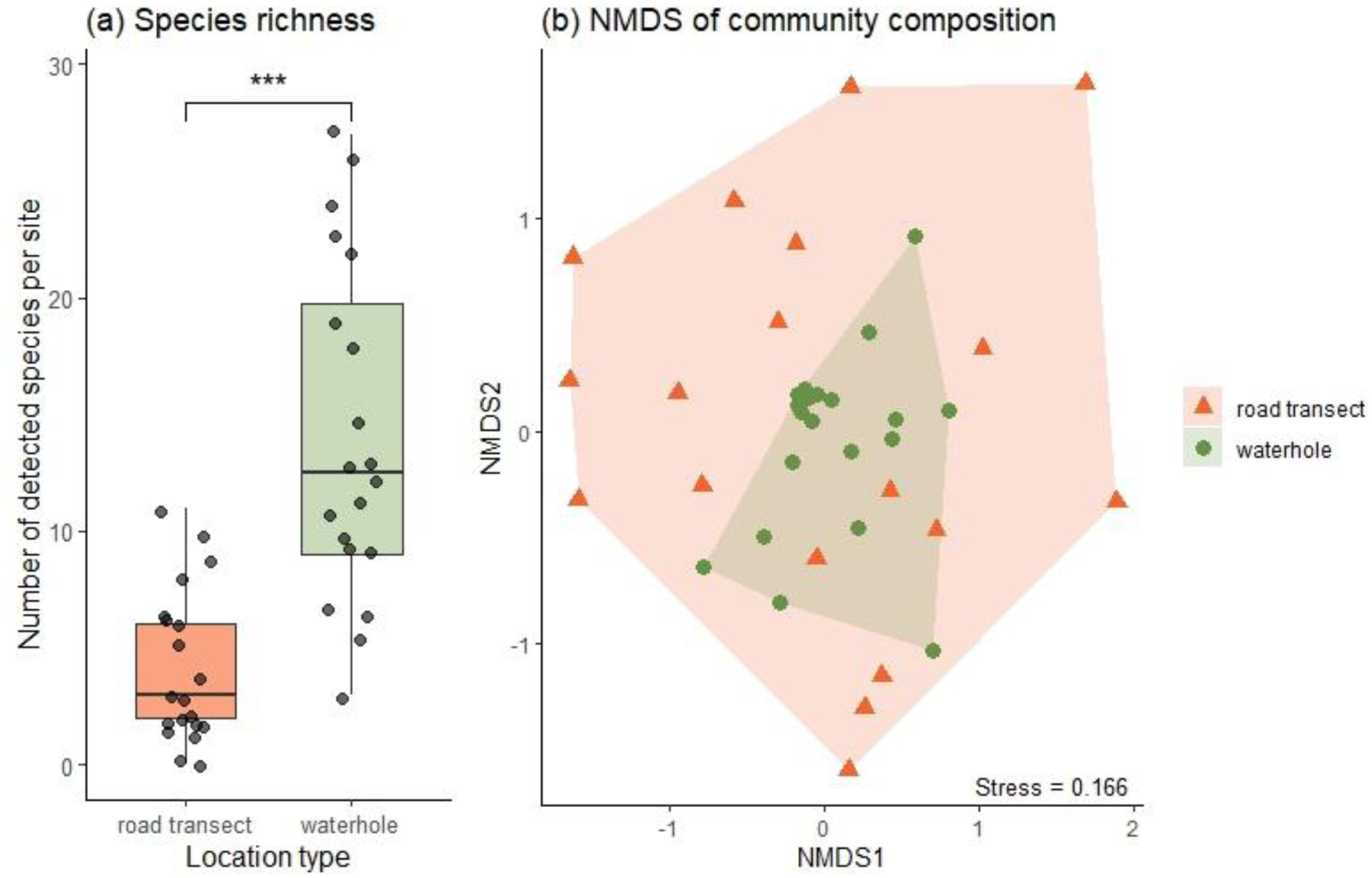
(a) Species richness per site at road transect and waterhole sampling sites. Boxes show the interquartile range and median whiskers indicate 1.5 × interquartile range, points represent individual sites. The horizontal bar and asterisks (***) indicate a highly significant difference in richness between location types based on the negative binomial GLM. (b) Non-metric multidimensional scaling (NMDS) ordination of community composition based on Jaccard dissimilarity of presence/absence data. Points represent sites, colored and shaped by location type. Polygons indicate convex hulls enclosing sites of the same type. The stress value is shown in the lower-right corner.

**Figure 4.**
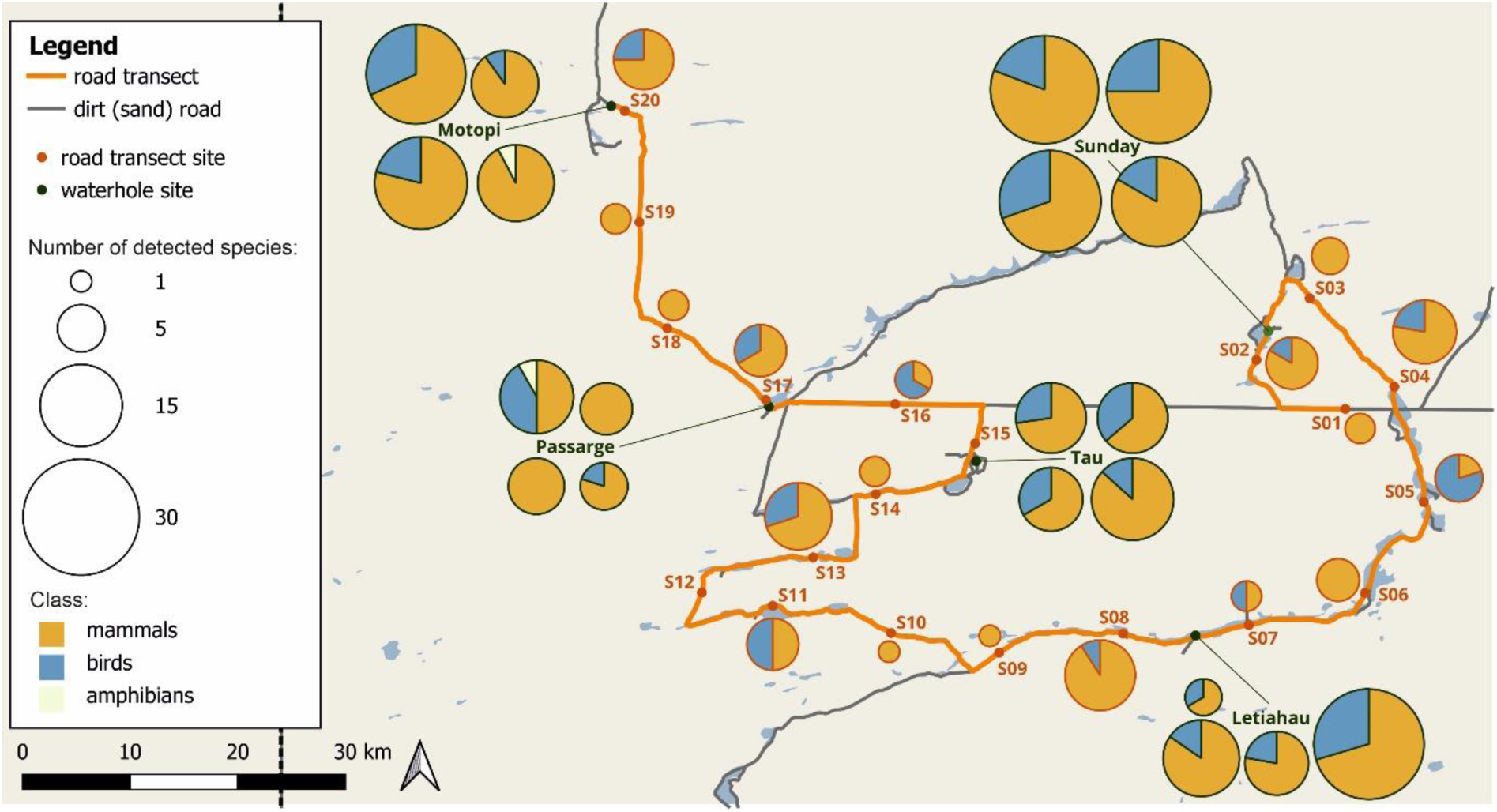
Detected biodiversity at each eDNA sampling site. Pie chart size indicates the total number of detected vertebrate species, while colored segments represent the relative contribution of each vertebrate class. Pie chart outlines indicate location type (green = waterhole; orange = road transect). Four sampling sites were surveyed per waterhole location, whereas one site was surveyed per road transect location.

**Table 3.**
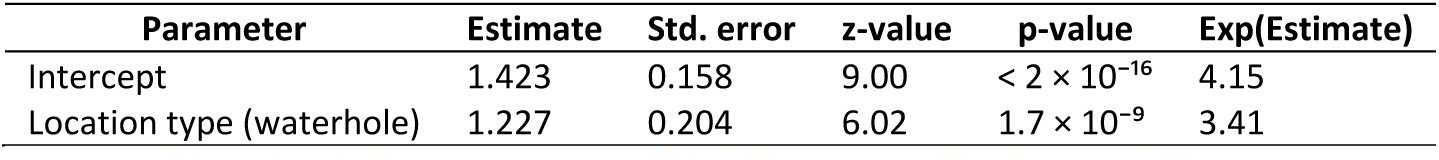
Negative binomial generalized linear model of species richness per site as a function of location type (reference category = road transect). Estimates are on the log scale; exponentiated coefficients give multiplicative effects on expected species richness.

### Isolation by distance

Mantel test showed no evidence for isolation by distance among road transect sites. Community dissimilarity (Jaccard distance) was not significantly related to driving distance between sites, as indicated by the Mantel test (Mantel r = 0.007, p = 0.454, 9,999 permutations, Figure S1).

### Comparison to documented vertebrate biodiversity in public databases

GBIF listed a total of 428 vertebrate species directly or indirectly observed within the CKGR (Table 4), most of which are birds (348), followed by mammals (52), reptiles (26) and amphibians (2). When comparing these numbers to the number of species of the respective classes detected by the eDNA survey, we achieve a detection rate of 73% for mammals and 100% of amphibians, whereas detection rates of bird and reptile species were substantially lower (Table 4). However, the lists of the detected species were not completely overlapping, with species detected in one of the two datasets on both sides. Notably, eight mammal species were detected by the eDNA survey that were not listed in the GBIF dataset. Four of these had “preserved specimen” records of European museum collections that fall within the reserve. However, as many of these records contained the exact same GPS coordinates, the reliability of these coordinates is to be doubted, as they may have simply served as a proxy using the centroid of the Botswana area (Table S2).

**Table 4.** Comparison of vertebrate diversity recorded in the GBIF database and detected with the eDNA survey within the study area. For each vertebrate class, the table reports the total number of species listed in GBIF, the number and percentage of those species detected by eDNA, the number of species shared between both datasets, and the numbers of species exclusive to GBIF or to the eDNA survey. Totals across all taxonomic classes are shown in the bottom row.

| class | number of species in GBIF database | species detected with eDNA survey | % detected in eDNA survey | species in both datasets | species exclusive in GBIF database | species exclusive in eDNA data |
| --- | --- | --- | --- | --- | --- | --- |
| Mammalia | 52 | 38 | 73% | 29 | 22 | 8 |
| Aves | 348 | 15 | 4% | 13 | 335 | 2 |
| Amphibia | 2 | 2 | 100% | 2 | 1 | 1 |
| Reptilia | 26 | 0 | 0% | 0 | 26 | 0 |
| **Total** | **428** | **55** | **13%** | **44** | **384** | **11** |

### Comparison to mammal biodiversity recorded by camera trapping

The camera-trapping survey yielded 31,048 images with mammal species. A total of 29 mammal taxa identified to species level were detected. The five most frequently detected species were springbok (*Antidorcas marsupialis*; 12,013 images), gemsbok (*Oryx gazella*; 7,487 images), giraffe (*Giraffa giraffa*; 2,503 images), black-backed jackal (*Lupulella mesomelas*; 1,748 images), and kudu (*Tragelaphus strepsiceros*; 1,656 images). Combined with the 38 mammal species detected by eDNA, a total of 43 mammal species were recorded across both monitoring approaches. Of these, 24 species (56%) were detected by both methods, 14 species (33%) were detected exclusively by eDNA, and five species (12%) were detected solely by camera trapping (Table 5). Camera trapping recorded four felid species not detected by eDNA, as well as one hyaenid species, whereas eDNA detected four murids not detected by camera trapping. Additional species detected by eDNA but missed by camera trapping included several antelope species, two mongoose species, and single species of shrew, pangolin, and bat.

**Table 5.**
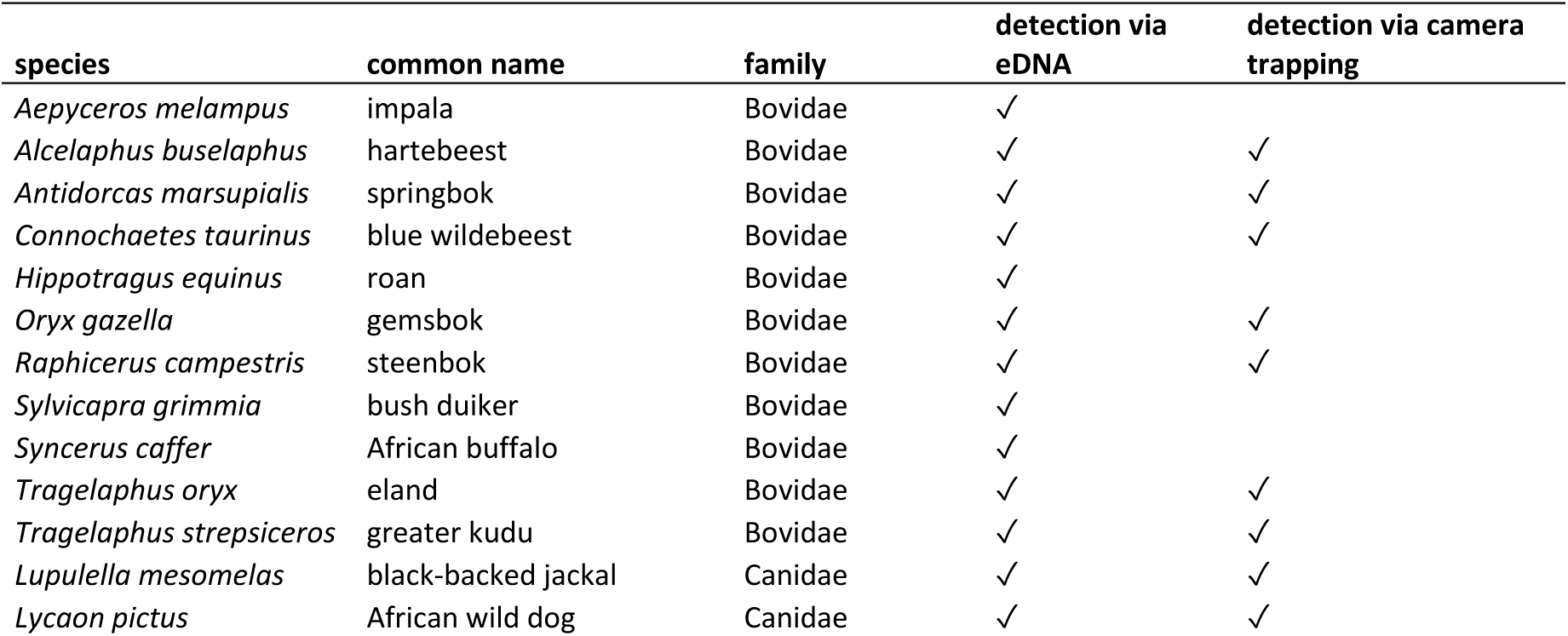
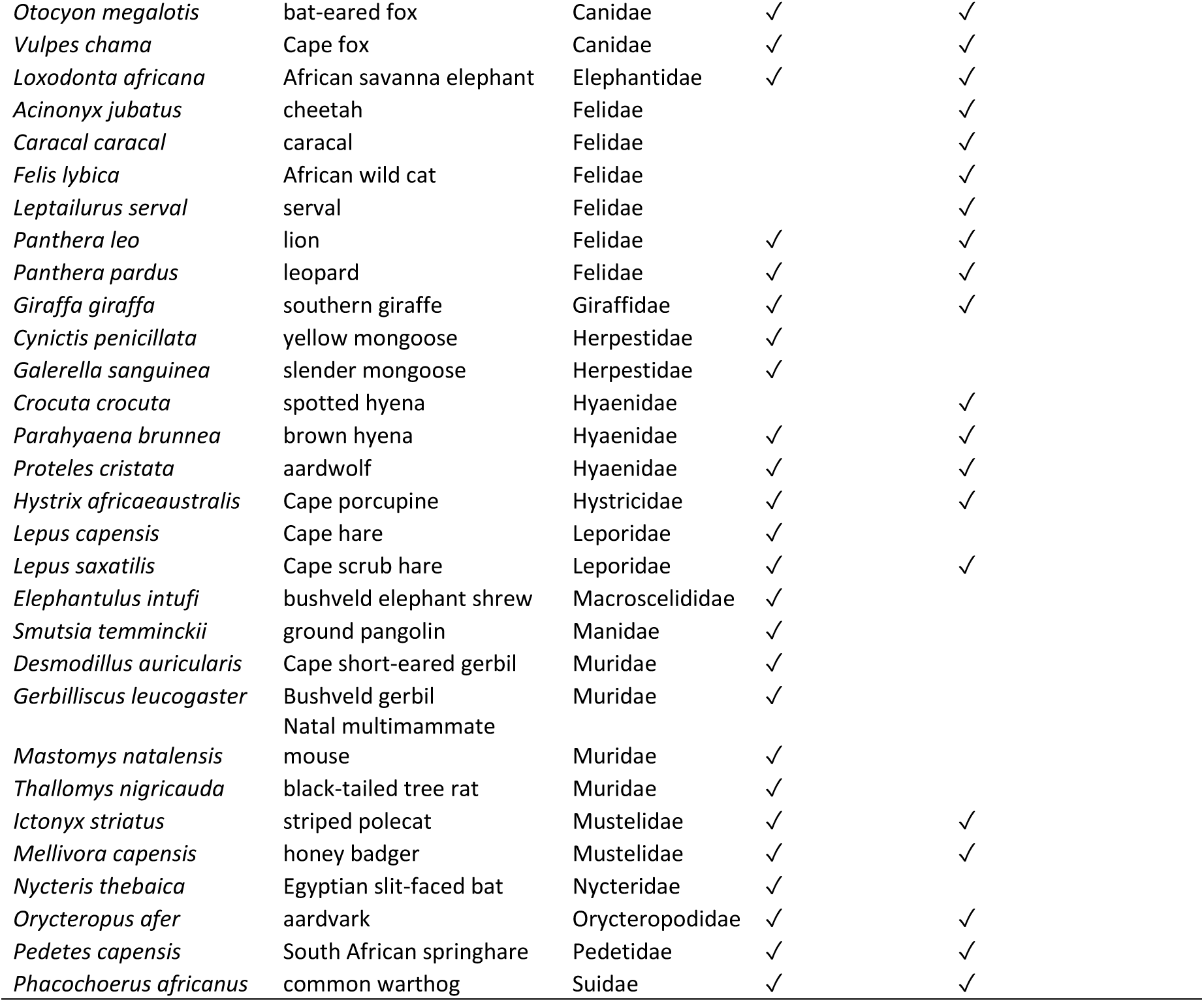
Mammal species detected by environmental DNA (eDNA), camera-trapping, or both methods. For each species, the scientific name, common name, and family are given, together with an indication of detection (✓) via eDNA and camera trapping.

There was a weak, positive association between the number of camera-trap image detections and the probability of eDNA detection (β ≈ 0.02, odds ratio ≈ 1.02 per additional image, p ≈ 0.08) across the entire dataset, meaning that species with more images tended to be more likely to be detected by eDNA. However, this effect was not statistically significant, indicating a weak trend rather than a strong, statistically robust relationship. Contrastingly, there was a statistically significant positive effect of the distance-weighted camera activity on eDNA detection probability (β = 0.092 ± 0.040, z = 2.28, p = 0.023). This indicates that species with greater camera trap activity near an eDNA sampling site were clearly more likely to be detected via eDNA.

## Discussion

Biodiversity loss is occurring at an unprecedented rate in the Anthropocene. The loss of large mammals is particularly severe due to habitat loss, overhunting, and climate change (Midgley and Thuiller 2011; Ripple et al. 2015). Some of their last remaining intact populations are found in remote and protected pockets of the world that are challenging to access and thus difficult to monitor. Here we demonstrate the feasibility of an eDNA-metabarcoding based approach to effectively capture mammal diversity in the remote CKGR in Botswana using an easy ‘scoop-up soil into a tube’ sampling protocol.

The applied sampling protocol could be carried out without the need for bulky equipment or cool-chain storage during field work and minimal single-use plastic production. No indication of contamination during field work (neither in laboratory workflow later) was observed in the negative controls, confirming a contamination-resistant approach for sampling terrestrial vertebrate eDNA in a remote arid environment. The quick sample collection (< 3 min per sample) and low material costs (∼ 1 $ per sample) make this approach also an attractive approach from an economical perspective.

Our eDNA survey revealed 55 vertebrate species within the CKGR, including 38 mammals, 15 birds and two amphibians overall. Two of the detected mammals (African buffalo *Syncerus caffer* and roan antelope *Hippotragus equinus*) do not typically occur in the CKGR (IUCN 2026) and generally prefer less arid habitats. However, a limited number of citizen-science observations have been recorded for both species within the CKGR. Their detections in our dataset may therefore indicate true presence at very low abundances, or alternatively, reflect secondary DNA input (e.g., via car tyres or scavenging birds).

In general, the waterhole-focused sampling approach yielded significantly higher species richness per sample, a greater total number of species detected, and more consistent species detections per sample than the road transect sampling. This pattern supports the hypothesis that eDNA of terrestrial vertebrates accumulates at these key aggregation points, consistent with previous descriptions from other semiarid and arid systems (Seeber et al. 2019; McDonald et al. 2023; Schenekar et al. 2026). The contrast was particularly pronounced for birds, with more than twice as many bird species detected at waterholes compared to road transects. In this densely vegetated, thorny bush landscape, wild mammals readily use roads, facilitating their detection along road transects (Abrahms et al. 2016; Fitzpatrick et al. 2024), whereas birds are not constrained to follow roads and likely shed much less eDNA onto the sampled road surfaces. Moreover, both amphibian species were detected only in proximity to waterholes, reflecting their strong dependence on aquatic habitats and underscoring the suitability of waterholes for detecting water-associated vertebrates via eDNA. Taken together, waterhole-focused eDNA sampling offers a more cost- and time-efficient approach to capture resident terrestrial vertebrate biodiversity in dry ecosystems. However, because waterholes represent critical resources and focal aggregation points for wildlife, particular care must be taken to (a) minimize disturbance to animals during sampling, and (b) implement rigorous safety protocols to reduce the risk of dangerous human–wildlife encounters associated with the high density and activity of wildlife at these sites.

The heterogeneous species composition across road transects (Fig. 3b), suggests that they may carry some signal of more localised species diversity. However, we found no evidence for isolation by distance along the road transect. Ecologically, this suggests that, at the spatial scale covered by our sampling, terrestrial vertebrate communities within the studied area of the CKGR are relatively homogeneous. This is consistent with the reserve’s open, sparsely infrastructured landscape, where many mammals have large home ranges and can move freely over tens of kilometres without strong natural barriers, whereas artificial barriers such as the perimeter fencing pose significant constraints on broader-scale movements (Dures et al. 2021; Sianga et al. 2024). From an eDNA perspective, this pattern may also indicate that road-sourced signals integrate activity from a broad catchment due to recurrent mixing and eDNA carryover caused by vehicles travelling along the road, potentially blurring fine-scale spatial structure in eDNA-based community patterns.

When benchmarked against the publicly available vertebrate records for the reserve, the sensitivity of soil eDNA differed markedly among taxonomic groups. Relative to the 52 mammal species listed for CKGR in GBIF, our survey detected 73% of the nominal mammal species richness derived from the GBIF database and 100% of the amphibian species richness. In contrast, bird and reptile detection rates were substantially lower. A large portion of the species detected exclusively by eDNA, but not previously listed for the CKGR in GBIF, were small rodents like murids, sengis and gerbils (*Gerbilliscus leucogaster, Desmodillus auricularis, Elephantulus intufi, Mastomys natalensis*), as well as cryptic and nocturnal species (aardvark – *Orycteropus afer* and pangolin – *Smutsia temminckii*). These taxa are hard to observe directly which makes them intrinsically less likely to be documented by citizen scientists, a major source of records in public occurrence databases (Stevenson et al. 2023). A substantial fraction of the bird species recorded for the CKGR in GBIF but not detected in our soil eDNA samples are either long-distance migrants (Palearctic or intra-African) or nomadic species whose occurrence in the reserve is strongly seasonal or tied to rainfall (Chittenden et al. 2017). Therefore, many of these species may simply not have been present in the system at the time of sampling, which was conducted towards the end of dry season.

The integrative analysis of eDNA and camera trapping data revealed clear biases of both monitoring approaches. Camera trapping performed well for large carnivores: it recorded four felids and one hyaenid species that were missed by eDNA. Given the make-up of species that were missed by eDNA sampling, there are several plausible explanations. Firstly, cheetah, caracal and African wildcat are not water-dependent and get the majority of their moisture requirements directly from their meals, and as smaller bodied felids, these species are likely to avoid the use of both roads and waterholes to avoid larger, dominant competitors like leopard and lion (Madden 2018; Davis et al. 2023; Morin et al. 2024; Nisi et al. 2026). While serval are conversely, associated with water, only one individual serval was detected at one site on the camera survey, and the species occurs here at the very edge of their known range (F. Weise pers .comm.). In addition, felids have repeatedly been reported to be difficult to detect via eDNA due to their behavioural ecology (Sales et al. 2020; Lyet et al. 2021) leading to relatively low eDNA shedding rates. Lastly, spotted hyena occur at relatively low densities across the landscape, as they generally do not do well in low rainfall systems (Jones et al. 2021).

Conversely, eDNA was particularly powerful for detecting small-bodied and cryptic taxa that are notoriously difficult to capture with camera traps. Many of these species are small or behaviourally cryptic, may not reliably trigger infrared sensors, use microhabitats that rarely intersect camera fields of view, and can be difficult to identify to species even when photographed. Yet they continuously shed DNA into the soil as they move and forage, making them accessible to eDNA metabarcoding.

Our analysis further showed that eDNA detection probability is positively associated with local activity as recorded by camera traps. Across the entire dataset, at the species level, there was a weak but positive trend whereby mammals with larger numbers of camera trap images tended to be more likely to be detected via eDNA. More importantly, when we accounted for the spatial distance between image recordings and eDNA sampling sites, there was a clear, statistically significant correlation between eDNA detection and camera trap–derived species activity. Species with higher camera trap activity near a given eDNA sampling point had a substantially higher probability of being detected via eDNA at that site. This supports a mechanistic link between eDNA signals and local mammal activity: where mammals are more frequently present and moving, they shed more DNA into the environment, and eDNA detection probabilities increase.

Taken together, these results underscore the complementarity of eDNA and camera trapping for terrestrial mammal monitoring for biodiversity conservation. Camera trapping delivers detailed behavioural and demographic information on medium- to large-bodied mammals, facilitates individual identification for some species, and provides direct evidence of presence and relative activity along specific trails. eDNA, by contrast, offers a rapid, low-effort snapshot of mammal community composition that is particularly sensitive to small and cryptic taxa, and can be implemented at many sites with minimal field infrastructure and no equipment that must be left in the field, so no revisits to sampling sites are required. It integrates presence over time windows of days to weeks, which can be advantageous for detecting species that only occasionally pass a given point, but may also blur fine-scale temporal resolution relative to camera-based detections.

Communities have an important role to play in conserving wildlife. Our findings highlight the rich and unique biodiversity of the CKGR, which is protected in part through the conservation efforts of the Government of Botswana and its people. Conservation approaches that meaningfully incorporate indigenous knowledge of the peoples and their perspectives have been proposed as both more equitable and more effective strategies for biodiversity conservation (Dawson et al. 2021). Meaningful participation of San communities in conservation initiatives and scientific research, including studies such as this one, could take a range of forms, such as assisting with sample collection in difficult-to-access areas, contributing observations and records of species presence, identifying ideal sampling sites to detect rarer species, affording opportunities to directly incorporate indigenous ecological knowledge into research and monitoring. This study highlights the simplicity and low contamination risk of the applied eDNA protocol, suggesting that basic training of indigenous peoples to collect eDNA samples in remote areas could add significant value to biodiversity monitoring in remote and difficult to access landscapes. Such participation could strengthen locally grounded knowledge, stewardship, and decision-making, while providing a foundation for more sustainable, community-based conservation.

## Conclusions

Our study shows that a simple, soil-based eDNA metabarcoding protocol can reliably capture vertebrate biodiversity in a large, remote semi-arid savanna, with particularly strong performance for mammals and clear advantages at animal aggregation points such as waterholes. Direct comparison with camera trapping revealed that eDNA and cameras sample complementary portions of the mammal community: cameras excel at large, wide-ranging carnivores, whereas eDNA is uniquely powerful for small-bodied and cryptic species, and eDNA detection probabilities increase with local camera-derived activity. Together, these findings highlight that soil eDNA provides a practical, scalable backbone for mammal biomonitoring in dry, data-poor protected areas, best used in combination with camera traps and other methods to achieve robust, community-level assessments.

## Acknowledgements

We would like to thank the Department of Environmental Protection / Ministry of Environmental and Tourism of the Government of the Republic of Botswana for providing the necessary permits for this research. The eDNA sample collection was performed under Research Permit ENT 8/36/4 LIX (106) and the research was approved by the Access and Benefit Sharing National Committee DEP 1/11/11 I (67). This research was funded in whole, or in part, by the Austrian Science Fund (FWF) [Grant DOI: 10.55776/35059-B]. For the purpose of Open Access, the author has applied a CC by public copyright licence to any Author-Accepted Manuscript (AAM) version arising from this submission. The authors acknowledge the financial support from the University of Graz and the Diversity & Global Connections Grant from the University of Bern.

