## Supplementary material for "Simple soil eDNA sampling reveals terrestrial mammal diversity in the Central Kalahari and highlights waterholes as sampling hotspots": Figure S1

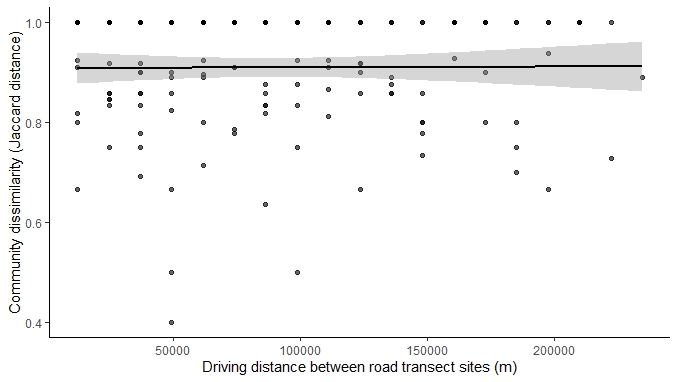

**Supplementary Figure** **S1** Relationship between geographic distance and community dissimilarity among road transect sites. Each point represents a pairwise comparison between sites, with the x-axis showing driving distance (m) and the y-axis showing Jaccard community dissimilarity (presence–absence data). The solid line and shaded band indicate the fitted linear regression and its 95% confidence interval.
