## Supporting Information 1 for "Simple soil eDNA sampling reveals terrestrial mammal diversity in the Central Kalahari and highlights waterholes as sampling hotspots": A_biodiversity comparison site type and location type.R

```
# Script to compare biodiversity detected at waterholes sites versus transect site and for creating Venn diagrams, boxplots and NMDS

# Set wd
setwd(dirname(rstudioapi::getActiveDocumentContext()$path))
# Load packages
library(dplyr) 
library(MASS) 
library(DHARMa)
library(vegan)  
library(ggplot2)
library(gridExtra) 
# Load csv


dat_by_site <- read.csv2("A_detections_by_site.csv", 
                             stringsAsFactors = FALSE) 

# Inspect the data
str(dat_by_site[, c("species_num", "loc_type")])  
summary(dat_by_site$species_num)  
table(dat_by_site$loc_type)  
dat_by_site$loc_type <- as.factor(dat_by_site$loc_type)  
hist(dat_by_site$species_num, breaks = 20)  

# glm 

dat_by_site$loc_type <- as.factor(dat_by_site$loc_type)  

m_pois <- glm(species_num ~ loc_type,  
              data   = dat_by_site,  
              family = poisson)  

summary(m_pois)  

# Test if loc_type has an effect:  
anova(m_pois, test = "Chisq")  

# Checking for overdispersion
overdispersion_stat <- sum(residuals(m_pois, type = "pearson")^2) / m_pois$df.residual  
overdispersion_stat  


# Overdispersion happening, switching to negative binomial

dat_by_site$loc_type <- as.factor(dat_by_site$loc_type)  

m_nb <- glm.nb(species_num ~ loc_type, data = dat_by_site)  

summary(m_nb)  
res <- simulateResiduals(m_nb, n = 1000)  
plot(res)  
testDispersion(res)  
testZeroInflation(res) 


anova(m_nb, test = "Chisq")  


# Venn diagrams

library(eulerr)  

# Load species list

species_list <- read.csv2(file = "A_Species detections list.csv")


  
# Make a presence absence matrix
species_pa <- species_list %>%  
  mutate(  
    present_waterhole = number.of.waterhole.locations  > 0,  
    present_transect  = number.of.road.transect.locations   > 0  
  )  

# Presence/absence categories  
only_waterhole <- species_pa %>%  
  filter(present_waterhole & !present_transect)  

only_transect <- species_pa %>%  
  filter(!present_waterhole & present_transect)  

both_site_types <- species_pa %>%  
  filter(present_waterhole & present_transect)  

# Counts for Euler diagram  
venn_data <- c(  
  "Waterhole"          = nrow(only_waterhole),  
  "Transect"           = nrow(only_transect),  
  "Waterhole&Transect" = nrow(both_site_types)  
)  

venn_data  


# Fit proportional Euler diagram  
fit <- euler(venn_data)  

palette_colors <- c("#c9dbba", "#faa381")  

Plot_waterhole_transect_species <- plot(  
  fit,  
  labels = list(  
    fontsize = 16,  
    position = "outside"  
  ),  
  quantities = list(  
    fontsize = 16  
  ),  
  edges = FALSE,  
  fills = palette_colors[1:2]  # first 2 colors for the two sets  
)  

Plot_waterhole_transect_species  


# Calculate alpha diversity (species richness/count ) for each loc_type

# Summary of richness by location type  
alpha_summary <- dat_by_site %>%  
  group_by(loc_type) %>%  
  summarise(  
    n_sites   = n(),  
    mean_S    = mean(species_num, na.rm = TRUE),  
    sd_S      = sd(species_num, na.rm = TRUE),  
    median_S  = median(species_num, na.rm = TRUE),  
    min_S     = min(species_num, na.rm = TRUE),  
    max_S     = max(species_num, na.rm = TRUE)  
  )  

alpha_summary  


######### Analysising beta diversity##########


# Defining species columns
species_cols <- 8:62  
sp.mat <- dat_by_site[, species_cols] 
# Keep row names as site IDs 
rownames(sp.mat) <- dat_by_site$site  
# Make sure species data are numeric 0/1  
sp.mat <- as.data.frame(lapply(sp.mat, as.numeric))  
# Grouping factor  
group <- dat_by_site$loc_type   


# Quick sanity checks  
dim(sp.mat)  
head(sp.mat[, 1:5])  
table(group)  
group <- factor(group)  

# Compute dissimilarity matrix.
# Identify rows with no species at all  
row_sums <- rowSums(sp.mat)  
row_sums  
empty_rows <- row_sums == 0  

# How many and which sites?  
sum(empty_rows)  
which(empty_rows)  
dat_by_site$site[empty_rows]  

# Remove them from species matrix and grouping factor  
sp.mat.nz <- sp.mat[!empty_rows, ]  
group.nz  <- droplevels(group[!empty_rows])  

# Compute Jaccard dissimilarity, binary = TRUE to interpret as presence/absence   
dist.jac <- vegdist(sp.mat.nz, method = "jaccard", binary = TRUE)  
dist.jac 


# PERMANOVA to see whether loc_type differs between loc_type

adonis_res <- adonis2(dist.jac ~ group.nz)  
adonis_res  
# Species differ between road and waterhole samples


# Boxplot of number of species per site by location type  
library(ggplot2)  

p_alpha_box <- ggplot(dat_by_site,  
                      aes(x = loc_type, y = species_num, fill = loc_type)) +  
  geom_boxplot(width = 0.6, outlier.shape = 21, outlier.fill = "white") +  
  geom_jitter(width = 0.15, alpha = 0.6, size = 2, color = "black") +  
  scale_fill_manual(values = c("road transect" = "#faa381",  
                               "waterhole"     = "#c9dbba")) +  
  labs(title = "(a) Species richness",
       x = "Location type",  
       y = "Number of detected species per site") +  
  theme_classic() +  
  theme(  
    legend.position = "none",  
    axis.title = element_text(size = 12),  
    axis.text  = element_text(size = 10)  
  ) +  
  ## ---- significance bar + stars ----  
# horizontal bar between the two groups  
geom_segment(aes(x = 1, xend = 2, y =  max(species_num) * 1.05,  
                 yend = max(species_num) * 1.05),  
             inherit.aes = FALSE) +  
  # small vertical ticks above each box  
  geom_segment(aes(x = 1, xend = 1,  
                   y = max(species_num) * 1.02,  
                   yend = max(species_num) * 1.05),  
               inherit.aes = FALSE) +  
  geom_segment(aes(x = 2, xend = 2,  
                   y = max(species_num) * 1.02,  
                   yend = max(species_num) * 1.05),  
               inherit.aes = FALSE) +  
  # text with PERMANOVA significance  
  annotate("text", x = 1.5, y = max(dat_by_site$species_num) * 1.08,  
           label = "***", size = 5)  

p_alpha_box   


# Make NMDS plot of species communities:
set.seed(123)  
nmds <- metaMDS(sp.mat.nz, distance = "jaccard", binary = TRUE, k = 2, trymax = 100)  


# Draw the plots with ggplot2

## 1. Extract site scores from the NMDS  
nmds_coords <- as.data.frame(scores(nmds, display = "sites"))  
nmds_coords$group <- group.nz   # add your factor  

## 2. Compute convex hulls for each group  
hull_data <- nmds_coords %>%  
  group_by(group) %>%  
  slice(chull(NMDS1, NMDS2)) %>%  
  ungroup()  

# 3. get the stress
stress_val <- nmds$stress 
## 4. Plot NMDS with convex hulls in ggplot  
# Site scores + group  
nmds_coords <- as.data.frame(scores(nmds, display = "sites"))  
nmds_coords$group <- group.nz  

# Convex hulls  
hull_data <- nmds_coords %>%  
  group_by(group) %>%  
  slice(chull(NMDS1, NMDS2)) %>%  
  ungroup()  

# Stress label  
stress_label <- paste0("Stress = ", round(nmds$stress, 3))  

plot_nmds_groups <- ggplot(nmds_coords,  
                           aes(x = NMDS1, y = NMDS2,  
                               color = group, shape = group)) +  
  geom_polygon(data = hull_data,  
               aes(x = NMDS1, y = NMDS2,  
                   fill = group, group = group),  
               alpha = 0.2,  
               inherit.aes = FALSE) +  
  geom_point(size = 3) +  
  annotate("text",  
           x = Inf, y = -Inf,  
           label = stress_label,  
           hjust = 1.1, vjust = -0.5,  
           size = 3.5) +  
  theme_classic() +  
  labs(title = "(b) NMDS of community composition",  
       x = "NMDS1",  
       y = "NMDS2") +  
  scale_color_manual(values = c("road transect" = "#F2652E",  
                                "waterhole"    = "#689147")) +  
  scale_fill_manual(values  = c("road transect" = "#F2652E",  
                                "waterhole"    = "#689147")) +  
  scale_shape_manual(values = c(17, 16)) +  
  theme(legend.title = element_blank())  

plot_nmds_groups  


grid.arrange(p_alpha_box,  plot_nmds_groups,
             ncol = 2,  
             widths = c(2, 4))
```
