## Supporting Information 1 for "Simple soil eDNA sampling reveals terrestrial mammal diversity in the Central Kalahari and highlights waterholes as sampling hotspots": B_isolation by distance.R

```
# Script to assess isolation by distance  

# Set wd  
setwd(dirname(rstudioapi::getActiveDocumentContext()$path))  

# Load packages  
library(dplyr)  
library(vegan)
library(ggplot2)  

# -----------------------------  
# 1. Load data  
# -----------------------------  
dat_by_site <- read.csv2("B_detections_by_site.csv",  
                         stringsAsFactors = FALSE)  

dat_road <- dat_by_site %>%  
  filter(loc_type == "road transect")  

dat_distances <- read.csv2("B_distance matrix.csv",  
                           stringsAsFactors = FALSE)  

# -----------------------------  
# 2. Define species columns and build community matrix  
# -----------------------------  
# Identify the first and last species columns by name:  
first_species_col <- which(names(dat_road) == "P_adspersus")  
last_species_col  <- which(names(dat_road) == "V_chama")  

comm_road <- dat_road[, first_species_col:last_species_col]  

# Make it presence/absence  
comm_road_pa <- comm_road  
comm_road_pa[comm_road_pa > 0] <- 1  

# Give community matrix rows the site IDs  
rownames(comm_road_pa) <- dat_road$site  

# -----------------------------  
# 3. Prepare distance matrix, align to road sites  
# -----------------------------  
# Extract road transect site IDs  
road_sites <- dat_road$site  

# Use first column "X" as row names  
rownames(dat_distances) <- dat_distances$X  
dat_distances$X <- NULL  # drop the column  

# Transform to matrix  
dist_mat <- as.matrix(dat_distances)  

# Keep only road-transect sites and order them consistently  
dist_road <- dist_mat[road_sites, road_sites]  

# Check that row/col names match the sites in dat_road  
all(rownames(dist_road) == dat_road$site)  

# -----------------------------  
# 4. Remove sites with no species detected  
# -----------------------------  
# Find rows (sites) with at least one species present  
non_empty <- rowSums(comm_road_pa) > 0  

# Which sites are empty?  
empty_sites <- dat_road$site[!non_empty]  

# Keep only non-empty sites in community data  
comm_road_pa_ne <- comm_road_pa[non_empty, ]  

# Also keep only non-empty sites in the metadata and distance matrix  
dat_road_ne  <- dat_road[non_empty, ]  
dist_road_ne <- dist_road[dat_road_ne$site, dat_road_ne$site]  

# -----------------------------  
# 5. Quick sanity checks  
# -----------------------------  
all(rownames(comm_road_pa_ne) == dat_road_ne$site)  
all(rownames(dist_road_ne)    == dat_road_ne$site)  

# -----------------------------  
# 6. Community dissimilarity: Jaccard  
# -----------------------------  
 

comm_dist_ne <- vegdist(comm_road_pa_ne,  
                        method = "jaccard",  
                        binary = TRUE)  

# Check that there are no NA distances  
any(is.na(comm_dist_ne))  

# -----------------------------  
# 7. Geographic distance as 'dist' object  
# -----------------------------  
geo_dist_ne <- as.dist(dist_road_ne)  

# Check that site labels align  
all(attr(comm_dist_ne, "Labels") == attr(geo_dist_ne, "Labels"))  

# -----------------------------  
# 8. Mantel test (Isolation by distance)  
# -----------------------------  
set.seed(123) 

mantel_res <- mantel(comm_dist_ne,  
                     geo_dist_ne,  
                     method = "pearson",  
                     permutations = 9999)  

mantel_res  


# -----------------------------  
# 9. Visualization  
# -----------------------------  


# Extract pairwise values from the 'dist' objects  
comm_vec <- as.vector(comm_dist_ne)  
geo_vec  <- as.vector(geo_dist_ne)  

df_ibd <- data.frame(  
  geo_dist  = geo_vec,  
  comm_dist = comm_vec  
)  


ggplot(df_ibd, aes(x = geo_dist, y = comm_dist)) +  
  geom_point(alpha = 0.6) +  
  geom_smooth(method = "lm", se = TRUE, color = "black") +  
  labs(x = "Driving distance between road transect sites (m)",  
       y = "Community dissimilarity (Jaccard distance)") +  
  theme_classic()  
```
