## Supporting Information 1 for "Simple soil eDNA sampling reveals terrestrial mammal diversity in the Central Kalahari and highlights waterholes as sampling hotspots": C_Camera_trapping_eDNA_comparison.R

```
# Script to compare eDNA and camera trapping data from CKGR surveys 2024

#SetWD
setwd(dirname(rstudioapi::getActiveDocumentContext()$path))

# Load required packages

library(lubridate)
library(dplyr)
library(readr)
library(tidyr)


# Import eDNA species list csv 
species_list_eDNA <- read_csv2("./C_eDNA_species_list.csv") 

# Keep only Mammalia in the 'class' variable
species_list_eDNA <- species_list_eDNA %>%
  filter(class == "Mammalia")

# Select only 'Species name' and 'Common name' and export as CSV
species_export <- species_list_eDNA %>%
  dplyr::select(`species name`, `common name`)

# Export eDNA mammal species names
write_csv2(
  species_export,
  "./CKGR_eDNA_Mammalia_species_common_names.csv")


# Import CT species list csv 
species_list_CT <- read_csv2(
  "C_CT_species_list_incl_common_names.csv",
  locale = locale(encoding = "UTF-8")
)

# Keep only rows where 'Common name' is not empty in species_list_CT
species_list_CT <- species_list_CT %>%
  filter(!is.na(`common name`) & `common name` != "")

############################### Comparing species lists of both detection methods ############################

# Make sure species names are character and unique per dataset
eDNA_species <- species_list_eDNA %>%
  mutate(`Species name` = as.character(`species name`),
         `Species name` = trimws(`species name`)) %>%
  distinct(`Species name`)

CT_species <- species_list_CT %>%
  mutate(`Species name` = as.character(`species name`),
         `Species name` = trimws(`species name`)) %>%
  distinct(`Species name`)

# Full join to see presence in each dataset
species_compare <- full_join(
  eDNA_species %>% mutate(in_eDNA = TRUE),
  CT_species   %>% mutate(in_CT   = TRUE),
  by = "Species name"
)

# Replace NA with FALSE for presence flags
species_compare <- species_compare %>%
  mutate(
    in_eDNA = if_else(is.na(in_eDNA), FALSE, in_eDNA),
    in_CT   = if_else(is.na(in_CT),   FALSE, in_CT)
  )

# Export eDNA CT comparison table
write_csv2(
  species_compare,
  "./mammal_species_list_eDNA_CT.csv")


# Species in both datasets

species_both <- species_compare %>%
  filter(in_eDNA & in_CT)

species_both_count <- nrow(species_both)

species_both_count  # number of species in both
species_both        # list of species in both


# Species in eDNA only
species_only_eDNA <- species_compare %>%
  filter(in_eDNA & !in_CT)

species_only_eDNA_count <- nrow(species_only_eDNA)

species_only_eDNA_count  # number of species only in eDNA
species_only_eDNA        # list of species only in eDNA


# Species only in CT
species_only_CT <- species_compare %>%
  filter(!in_eDNA & in_CT)

species_only_CT_count <- nrow(species_only_CT)

species_only_CT_count  # number of species only in CT
species_only_CT        # list of species only in CT


####################### Logistic regression to see whether more CT image records increase the chance of detection via eDNA ###########
# 1) Extract CT detection counts per species
ct_counts <- species_list_CT %>%
  mutate(`species name` = trimws(as.character(`species name`))) %>%
  dplyr::select(`species name`, `number of detections total`) %>%
  distinct()

# 2) Merge CT counts into species_compare
#    (Note: species_compare has "Species name")
species_compare_counts <- species_compare %>%
  mutate(`species name` = trimws(as.character(`Species name`))) %>%
  left_join(ct_counts, by = "species name")

# 3) Restrict to species that occur in CT (have non-missing counts)
species_compare_ct <- species_compare_counts %>%
  filter(!is.na(`number of detections total`))

# 4) Convert eDNA detection to numeric (0/1) for logistic regression
species_compare_ct <- species_compare_ct %>%
  mutate(
    eDNA_detected = if_else(in_eDNA, 1L, 0L)
  )

# 5) Fit logistic regression:
#    eDNA_detected (binary) ~ number of CT detections (continuous)
model_eDNA_CT <- glm(
  eDNA_detected ~ `number of detections total`,
  data = species_compare_ct,
  family = binomial(link = "logit")
)

# 6) Inspect model results
summary(model_eDNA_CT)

# 7) Odds ratio for 'number of detections total'
odds_ratio <- exp(coef(model_eDNA_CT)["`number of detections total`"])
odds_ratio

# 8) 95% confidence interval for the odds ratio
confint_or <- exp(confint(model_eDNA_CT)["`number of detections total`", ])
confint_or
```
