## Supporting Information 1 for "Simple soil eDNA sampling reveals terrestrial mammal diversity in the Central Kalahari and highlights waterholes as sampling hotspots": D_spatial glmm CT on eDNA.R

```
# Script of spatially-aware modelling of the effect of CT recordings on eDNA detections

# by Tamara Schenekar

#SetWD
setwd(dirname(rstudioapi::getActiveDocumentContext()$path))

# Load required packages
library(tidyverse)
library(lubridate)
library(sf) 
library(lme4)
library(stringr)

## 1) Read and convert eDNA data
edna_raw <- read_csv2("Mammal_eDNA_detections_by_site.csv")

edna_clean <- edna_raw %>%
  ## Standardize column names for convenience
  rename(
    edna_site   = `Sampling site`,
    edna_lon_raw = Longitude,
    edna_lat_raw = Latitude,
    edna_time_raw = Timestamp
  ) %>%
  ## Replace comma with dot and convert to numeric
  mutate(
    edna_lon = as.numeric(str_replace(edna_lon_raw, ",", ".")),
    edna_lat = as.numeric(str_replace(edna_lat_raw, ",", "."))
  ) %>%
  ## Parse the timestamp; format appears to be "dd.mm.yyyy HH:MM"
  mutate(
    edna_time = dmy_hm(edna_time_raw)
  )


## 2) Read camera data
ct_raw   <- read_csv2("CT_data_EastWestsurvey_combined.csv")

ct_clean <- ct_raw %>%
  rename(
    survey     = `Survey Name`,
    ct_site    = `Site Name`,
    ct_lat_raw = Latitude,
    ct_lon_raw = Longitude,
    ct_species = Species,
    ct_time_raw = Timestamp,
    image_url  = `Image URL`
  ) %>%
  mutate(
    ct_lon = as.numeric(str_replace(ct_lon_raw, ",", ".")),
    ct_lat = as.numeric(str_replace(ct_lat_raw, ",", ".")),
    ct_time = dmy_hm(ct_time_raw)
  )

# Convert eDNA data from wide to long format

edna_long <- edna_clean %>%
  pivot_longer(
    cols = c(
      "Aepyceros melampus",
      "Alcelaphus buselaphus",
      "Antidorcas marsupialis",
      "Canis mesomelas",
      "Connochaetes taurinus",
      "Cynictis penicillata",
      "Desmodillus auricularis",
      "Elephantulus intufi",
      "Galerella sanguinea",
      "Gerbilliscus leucogaster",
      "Giraffa giraffa",
      "Hippotragus equinus",
      "Hystrix africaeaustralis",
      "Ictonyx striatus",
      "Lepus capensis",
      "Lepus saxatilis",
      "Loxodonta africana",
      "Lycaon pictus",
      "Mastomys natalensis",
      "Mellivora capensis",
      "Nycteris thebaica",
      "Orycteropus afer",
      "Oryx gazella",
      "Otocyon megalotis",
      "Panthera leo",
      "Panthera pardus",
      "Parahyaena brunnea",
      "Pedetes capensis",
      "Phacochoerus africanus",
      "Proteles cristata",
      "Raphicerus campestris",
      "Smutsia temminckii",
      "Sylvicapra grimmia",
      "Syncerus caffer",
      "Thallomys nigricauda",
      "Tragelaphus oryx",
      "Tragelaphus strepsiceros",
      "Vulpes chama",
      "Acinonyx jubatus",
      "Caracal caracal",
      "Felis lybica",
      "Leptailurus serval",
      "Crocuta crocuta"
    ),
    names_to   = "species",
    values_to  = "edna_detect"
  ) %>%
  mutate(
    species     = as.factor(species),
    edna_detect = as.integer(edna_detect)
  )


# Summarise images per camera site x species
ct_site_species <- ct_clean %>%
  group_by(ct_site, ct_lon, ct_lat, ct_species) %>%
  summarise(
    n_images = n(),         # number of images for this species at this camera
    .groups  = "drop"
  )


# Restrict species set only to the ones appearing in both eDNA and CT dataset
species_edna <- levels(edna_long$species)
species_ct   <- unique(ct_site_species$ct_species)

length(species_edna)
length(species_ct)

# Intersection: species that appear in BOTH datasets
species_overlap <- intersect(species_edna, species_ct)
length(species_overlap)
species_overlap

ct_site_species_overlap <- ct_site_species %>%
  filter(ct_species %in% species_overlap)


##### Calculating the distances between sites #####################
# 5.1 create spatial objects for eDNA sites and camera sites
## Unique eDNA sites
edna_sites <- edna_long %>%
  distinct(edna_site, edna_lon, edna_lat) %>%
  st_as_sf(coords = c("edna_lon", "edna_lat"), crs = 4326)  # WGS84

## Unique camera sites
ct_sites <- ct_site_species_overlap %>%
  distinct(ct_site, ct_lon, ct_lat) %>%
  st_as_sf(coords = c("ct_lon", "ct_lat"), crs = 4326)      # WGS84

edna_sites
ct_sites
nrow(edna_sites)  # should be 40
nrow(ct_sites)    # number of camera sites


# 5.2 Project to a metric CRS and compute distance matrix
edna_sites_utm <- st_transform(edna_sites, crs = 32734)
ct_sites_utm   <- st_transform(ct_sites,   crs = 32734)

## Distance matrix: eDNA sites (rows) × camera sites (columns)
dist_matrix <- st_distance(edna_sites_utm, ct_sites_utm)  # units: meters
dim(dist_matrix)


# Convert distance matrix to da tidy frame
## Get site IDs back from sf objects
edna_sites_df <- edna_sites_utm %>% st_drop_geometry()
ct_sites_df   <- ct_sites_utm   %>% st_drop_geometry()

## Turn distance matrix into a data frame and long format
dist_df <- as.data.frame(dist_matrix)

# Name columns as ct_col_1..N
colnames(dist_df) <- paste0("ct_col_", seq_len(ncol(dist_df)))

dist_df <- dist_df %>%
  mutate(edna_site = edna_sites_df$edna_site) %>%
  pivot_longer(
    cols      = starts_with("ct_col_"),
    names_to  = "ct_col",
    values_to = "distance_m"
  ) %>%
  mutate(
    ct_index = as.integer(str_remove(ct_col, "ct_col_")),
    ct_site  = ct_sites_df$ct_site[ct_index]
  ) %>%
  select(edna_site, ct_site, distance_m)


# 6. Combine distances with camera species activity
edna_camera_combined <- dist_df %>%
  inner_join(ct_site_species_overlap, by = "ct_site")

# Convert distance_m (units) to numeric meters
edna_camera_combined <- edna_camera_combined %>%
  mutate(distance_m_num = as.numeric(distance_m))

glimpse(edna_camera_combined)
summary(edna_camera_combined$distance_m_num)

# Compute weights and weighted images using the numeric distance
lambda <- 5000  # distance decay scale in meters

edna_camera_combined <- edna_camera_combined %>%
  mutate(
    weight          = exp(-distance_m_num / lambda),
    weighted_images = weight * n_images
  )

glimpse(edna_camera_combined)
summary(edna_camera_combined$weighted_images)


# Create a Camera index that summarises all the weighted images
camera_index <- edna_camera_combined %>%
  group_by(edna_site, ct_species) %>%
  summarise(
    cam_index = sum(weighted_images),
    .groups   = "drop"
  )


# Attach the camera index to eDNA long

# Restricht to only species detected

# species that have at least one eDNA detection
species_with_detection <- edna_long %>%
  group_by(species) %>%
  summarise(any_detection = any(edna_detect > 0), .groups = "drop") %>%
  filter(any_detection) %>%
  select(species)

edna_long_detected <- edna_long %>%
  semi_join(species_with_detection, by = "species")
# Attach/ join
edna_with_cam <- edna_long_detected %>%
  left_join(
    camera_index,
    by = c("edna_site" = "edna_site",
           "species"  = "ct_species")
  ) %>%
  mutate(
    cam_index = replace_na(cam_index, 0)  # species/site with no camera info get 0
  )

glimpse(edna_with_cam)
summary(edna_with_cam$cam_index)


# Fit the glmm model


mod1 <- glmer(
  edna_detect ~ cam_index + (1 | edna_site) + (1 | species),
  data   = edna_with_cam,
  family = binomial(link = "logit")
)

summary(mod1)


# Rescaling of variables suggested
edna_with_cam <- edna_with_cam %>%
  mutate(cam_index_100 = cam_index / 100)

summary(edna_with_cam$cam_index_100)


mod2 <- glmer(
  edna_detect ~ cam_index_100 + (1 | edna_site) + (1 | species),
  data   = edna_with_cam,
  family = binomial(link = "logit")
)

summary(mod2)

### camera index has a significant effect!


# Compare model to 0 model
mod0 <- glmer(
  edna_detect ~ 1 + (1 | edna_site) + (1 | species),
  data   = edna_with_cam,
  family = binomial(link = "logit")
)

anova(mod0, mod2)
```
